# Molecular unbalances between striosome and matrix compartments characterize the pathogenesis of Huntington’s disease model mouse

**DOI:** 10.1101/2025.07.20.665815

**Authors:** Ryoma Morigaki, Tomoko Yoshida, Joji Fujikawa, Jill R. Crittenden, Ann M. Graybiel

**Author notes:** Correspondence: Ann M. Graybiel; 43 Vassar Street 46-6133, Cambridge, MA 02139.

## Abstract

The pathogenesis of Huntington’s disease is still incompletely understood, despite the remarkable advances in identifying the molecular effects of the *Htt* mutation in this disease. When we focus on movement disorders, clinical studies offer us some hints about this issue. Human studies employing positron emission tomography have identified a reduction in phosphodiesterase 10A (PDE10A) as the earliest event in the brain of patients with Huntington’s disease, which occurs about 25 years before symptom onset. A *PDE10A* mutation is also known to cause childhood-onset chorea. *GNAL* encodes the olfactory type G-protein α subunit (Gα_olf_), strongly expressed in the striatum, and its mutation causes familial dystonia. PDE10A and Gα_olf_ are both critical regulators of cyclic AMP and are abundant in striatal spiny projection neurons. These findings suggest that maintaining cyclic AMP levels in the striatum might be an essential target for the pathogenesis of movement disorders such as chorea and dystonia. Why and how these changes in the striatum cause movement disorder are still a mystery. Here we suggest that a key might be evaluating these messenger systems in light of the circuit-level compartmental organization of the striatum, in which there is particular vulnerability of the striosome compartment. We developed machine learning algorithms to define with high precision and reproducibility the borders of striosomes in the brains of q175 Huntington’s disease model mice from 3-12 months of age. We demonstrate that multiple molecules including Gα_olf_, PDE10A, dopamine D1 and D2 receptors, adenosine 2A receptors, and mu-opioid receptors differentially change their expression patterns in striosomes across ages by comparison with their expression patterns in the matrix compartment. An early and pronounced differential vulnerability of striosomes has been demonstrated in studies of post-mortem Huntington’s brains. Our findings here, mapping the molecular distributions across age in a widely studied mouse model of Huntington’s disease, may help to pinpoint the pathogenic mechanisms of Huntington’s disease by demonstrating the differential molecular changes in the striosome compartment.

## Introduction

The differential neurodegeneration of the striosome and matrix compartments has been found in post-mortem neuropathologic studies of the striatum in patients with Huntington’s disease (HD) [1–12]. Further, abnormalities in gene regulation in the striosome-matrix axis or organization have also been found by single-nucleus RNA sequencing (snRNA-seq), accompanying the well-known decline in the spiny projection neurons (SPNs) expressing dopamine D2 receptors [13]. Examination of a Grade 1 case has suggested that SPNs of striosomes are likely the first SPNs to die, and alignment of early striosome-matrix imbalance with the early appearance of pre-manifest symptoms has been reported. The precise process of molecular alterations in these compartments that cause neuronal death and aberrant brain function in patients with HD is largely unknown. Hindering these studies is the small diameter of most of the striosome components (∼0.5-1 mm in the human) and their labyrinthine distribution within the much larger matrix compartment. These constraints so far made imaging of the striosomal system impossible or, at best, problematic. Nevertheless, a strong and reliable clue from imaging the striatum in HD patients has been derived from positron emission tomography (PET) studies [14,15]. These PET studies have identified a reduction in striatal phosphodiesterase 10A (PDE10A) as the earliest disease marker in premanifest HD patients. A decline in PDE10 precedes the decreases of dopamine D1 (Drd1) and dopamine D2 (Drd2) receptors [14,16], and PDE10A expression is reduced by 25–35% approximately 25 years before symptomatic onset. This reduction is mostly restricted to the dorsal sensorimotor striatum, and the decrease in expression was reported to be correlated with disease burden and severity [14,16]. PDE10A is highly expressed in SPNs, in which it regulates cyclic AMP (cAMP) and cyclic GMP (cGMP). It has an approximately 20-fold higher affinity for cAMP than for cGMP; it hydrolyzes both, which eventually produces inhibitory effects on cAMP/cGMP [17–19]. Intriguingly, heterozygous as well as bi-allelic mutations in *PDE10A* lead to childhood-onset chorea [20–22]. This clinical evidence suggests that PDE10A deficiency in the striatum, leading to dysregulation of cAMP levels in the striatum, might be a critical element accounting for the generation of movement disorders [23].

A second lead from the clinic concerns the olfactory type G-protein α subunit (Gα_olf_), a G protein related to movement disorders by dysregulating cAMP levels in the striatum. Mutation of the *GNAL* gene, which encodes Gα_olf_, is implicated as a cause of hereditary dystonia [24–32]. Gα_olf_ is highly concentrated in the striatum, where it couples positively with Drd1 and adenosine 2A receptors (A2A) in SPNs. When these receptors are stimulated, Gα_olf_ works to increase intracellular cAMP levels in SPNs [33,34]. Given that *PDE10A* and *GNAL* mutations critically affect the cAMP level in the striatum, one hypothesis is that several molecules expressed in SPNs might alter their expression to maintain the cAMP levels, resulting in unbalanced signaling transmission and movement dysfunction [23]. Of note, Gα_olf_ is expressed predominantly in the striosome compartment of the striatum [35,36]. The interaction between abnormalities in the striosome compartment and hyperkinetic movement disorders has already been suggested [37–39]. Differences in striosome and matrix involvement have been implied in the hyperkinetic movement disorder X-linked dystonia-parkinsonism (XDP; DYT/PARK-TAF1; Lubag) [40,41] and levodopa-induced dyskinesia [36,42,43], as well as in Huntington’s disease [5,7–9,11,12]. These correlations are consistent with the possibility, posed here, that imbalanced expression of molecules to maintain the cAMP levels in the striosome compartment might be one key factor in inducing some classes of movement disorder.

Given the lack of human brain imaging to visualize striosomes, we turned to immunohistochemical staining, a traditional method but still one of the best methods to define the striosome compartment. We have recently reported that mu-opioid receptor 1 (MOR1) in the striatum, mainly starting in the striosome compartment, is strikingly and progressively upregulated across ages in the Q175 knock-in (Q175KI) mouse model of HD [44]. Here, we undertook further immunohistochemical staining using Q175KI mice across three ages (3-, 6-, and 12-month-old). We found that Drd1, Drd2, A2A, PDE10A, and Gα_olf_ significantly decreased, especially in striosomes in the Q175KI mice compared to wildtype (WT) mice. This mapping leaves MOR1 as the sole upregulated receptor of this broad group. The spatio-temporal alteration of these molecules, although here documented only in a mouse HD model, could yield new insight into the pathogenesis of HD by demonstrating visible evidence of abnormalities in cAMP-related molecules, and the differential expression of these abnormalities in striosomes.

## Material and methods

### Animals

All experimental protocols involving work with mice were approved by the Committee on Animal Care at the Massachusetts Institute of Technology. For the primary analysis, twenty-four mice, four male and four female heterozygous Q175KI mice, and matching numbers for WT littermates for each of three ages (3, 6, and 12 months postnatal). Additionally, we used six mice each at 2, 6, and 19 months of age. All mice were obtained from Jackson Laboratory (Bar Harbor, ME, USA) or our laboratory colony at MIT.

### Immunizing peptide-blocking assay

The immunizing peptide-blocking assay for rabbit polyclonal PDE10A (Creative Diagnostics, Shirley, NY) was performed as described previously [44].

### Immunohistochemistry

Immunohistochemistry was performed as described previously [38,44,45]. In brief, 20-μm thick sections were cut on a sliding microtome, and free-floating sections were immunohistochemically stained. After a step for blocking endogenous peroxidase activity, sections were incubated in phosphate buffered saline (PBS) containing 3% bovine serum albumin (BSA) for 60 min and then were incubated in the chosen antibodies for 15 h (Supplementary Table 1). Bound antibody was detected by use of anti-rabbit (Invitrogen, Carlsberg, CA), anti-goat (Vector, Burlingame, CA), or anti-rat (Nichirei, Tokyo, JP) IgG polymer secondary antibodies, and bound peroxidase was detected by 3,3’-diamonobenzidine (DAB) (Vector, Burlingame, CA) or the TSA-system (fluorescein or Cy5) (Perkin Elmer, Shelton, CT). For dual antigen detection of fluorescent reagents, sections were incubated in 0.1 M glycine-HCl (pH 2.2) for 30 min after the first antigen procedures to remove the bound antibody. After several rinses in PBS, sections were incubated for 15 h in PBS containing 3% BSA and rabbit anti-MOR1. Bound antibody was detected the same as the first antibody. The sections from WT and Q175KI mice of the same ages for each target molecule were stained simultaneously.

### Immunohistochemical detection of PDE10A in monkeys’ brain

Studies on striatal sections from Macaca mulatta monkeys were conducted on monkeys from the laboratory collection of monkeys treated according to the Guide for Care and Use of Laboratory Animals of the United States National Institutes of Health and with the approval of the Committee on Animal Care at the Massachusetts Institute of Technology. Two monkeys were perfused through the heart with 0.9% saline, then 4% paraformaldehyde in a 0.1M phosphate-buffered 5% sucrose solution, and a post-wash with phosphate-buffered sucrose under deep barbiturate anesthesia (50 mg/kg, i.p.). The brains were soaked 3 days in 20% glycerol and cut coronally at 40 µm on a freezing microtome. Immunostaining was performed by the free-floating method, as for sections from mice, except that PBTA method was used for anti-PDE10A antibody visualization [46]. In brief, after incubation with anti-PDE10A antibody (1:100,000) for two days and several rinses with PBS, sections were incubated with anti-rabbit polymer-staining reagent (Invitrogen, Carlsberg, CA) for 30 min. After several rinses in PBS, the sections were processed for TSA with the TSA Biotin System (Perkin Elmer). Sections were incubated in the biotinyl tyramide amplification reagent (1:100 using 1×Plus Amplification Diluent) for 15 min. After several rinses in PBS, the sections were incubated with the avidin-biotin-peroxidase complex (ABC) reagent (Vector, Burlingame, CA) for 30 min. The bound peroxidase was visualized by incubating the sections with a solution containing 3,3’-diaminobenzidine with nickel (Vector) and 0.01% H_2_O_2_ for 2 min. For dual antigen detection, sections were processed the same as mice except using mouse anti-K-chip antibody (1:2,000; NeuroMab, Cambridge, MA) for second primary antibody and anti-mouse polymer secondary antibody.

### Striosomal segmentation using deep learning by a fully convolutional residual neural network U-Net

To perform the segmentation of striatal compartments automatically and objectively, we developed an algorithm using deep learning by a fully convolutional residual neural network U-Net [47]. We randomly selected 176 MOR1 fluorescent images from 2-, 3-, 6-, 9-, 12- and 19-month-old WT and Q175KI mice as training images (12,000×9,000 pixels). Firstly, two authors (RM and JF) manually defined all of the striosomes visible, and then fibers and vessels, with the assistance of Otsu’s method that performed automatic image thresholding [48]. The background intensities were set on the slide grass outside the tissue. Then, the dorsal striatum was divided into 2376 training images (512×512 pixels for each image). We tested two methods, augmentation with vertical flip or without augmentation for each image. The evaluation was performed using randomly selected 8 images (total 24 images, 48 striatum) from 3-, 6-, and 12-month-old WT and Q175KI mice. The batch size of the training set was 32 and the total epoch was 160. The training rate was set as 0.85.

### Digital images and densitometry

Microscopic images were captured with a TissueFAXS whole-slide scanning system equipped with a Zeiss Axio Imager Z2 fluorescence microscope and TissueFAXS image acquisition software as described before [43]. The acquired images were processed and analyzed with MATLAB software. The optical density of each molecule’s immunolabeling was entered as gray levels on non-colored digital images. Each molecule’s labeling was measured in the striosomal and matrix subfields of the striatum demarcated by rabbit monoclonal anti-MOR1 antibody for each mouse (n = 8, each with both hemispheres, yielding 16 caudoputamen samples) across each age (3, 6, and 12 months). The levels of the caudoputamen, coronal sections from +0.98 to +1.34 mm rostral to the bregma [49] were selected for each WT and Q175KI mouse. We arranged 24 serial sections (20 µm-thick) in a 96-well plate for each mouse and selected the brain region of these coordinates as nearly identical as possible for immunohistochemical staining of each molecule. Four segments in the caudoputamen (DM: dorsomedial, DL: dorsolateral, VM: ventromedial, and VL: ventrolateral) were manually defined [43]. We defined “WH” as the whole area of the caudoputamen, which was the sum of these four segments. Then the MOR1-immunopositive regions were objectively defined by the deep learning using U-net architecture. The MOR1-immunopositive striosomal regions were superimposed on the double-stained immunohistochemical images of each molecule. The densitometry analyses were performed for the striosome and matrix compartments, excluding fiber bundles. Segments exhibiting poor staining or tissue loss were excluded from subsequent analyses following visual inspection of the histological images. Each segment’s relative intensity (RI) was calculated as a ratio of mean optical intensity in Q175KI mice relative to the mean for WT for of each segment and age:

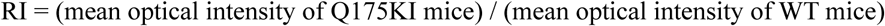

The relative ratio of striosome and matrix intensities, measured as an index of striosome-to-matrix predominance (ISMP), was calculated in each segment of the same animals [50,44]:

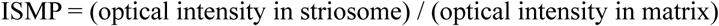

### Statistical analysis

All quantitative data were expressed as means ± SEM values. The Kruskal-Wallis tests followed by pairwise Mann-Whitney U-tests with Bonferroni corrections were used. *P*-values less than 0.05 were considered statistically significant.

## Results

### 1. Striosomal segmentation using deep learning by a fully convolutional residual neural network U-Net

The performance comparison of the two methods is shown in Supplementary Table 2. Augmentation with a vertical flip is better than without augmentation. This method was used for subsequent experiments, and representative images are shown in Supplementary Figure 1A-C. The model was stable with the lowest cross-entropy loss at 69 epochs by vertical flip without augmentation (accuracy and loss test, 0.97965556 and 0.05494046, respectively) and 44 epochs by vertical flip with augmentation (accuracy and loss test, 0.97895205 and 0.055600364, respectively).

### 2. MOR1 is upregulated in the striatum of the Q175KI mice

Confirming our earlier finding [44], MOR1 immunoreactivity was upregulated in the striatum of Q175KI mice compared to littermate WT mouse striatum (Figure 1). The upregulation was predominant in the striosome compartment, especially at 12-month-old. Compared to WT mice, Q175KI mice showed the increased intensity of MOR1 labeling in the striosome compartment at 3 and 6 months in the DL, and at 12 months in the WH, DM, DL, and VL segments. In the matrix compartment, the increase was limited to the 12-month DL segment. The ISMP ratios increased in 3- and 6-month DL, and 12-month WH, DM, DL, and VL segments (Figure 2A). These results indicated that MOR1 expression in striosomes increased in the DL segment as early as 3 months of age, and at the entire dorsal striatum except VM segment at 12-month-old. MOR1 expression in the matrix compartment began to increase at 12-month-old, as we also observed earlier [2]. This confirmation gives strength to this remarkable upregulation of the MOR1 labeling. As shown in Figure 1, MOR1 was the only molecule of those tested that progressively upregulated its expression in Q175KI mice.

**Figure 1:**
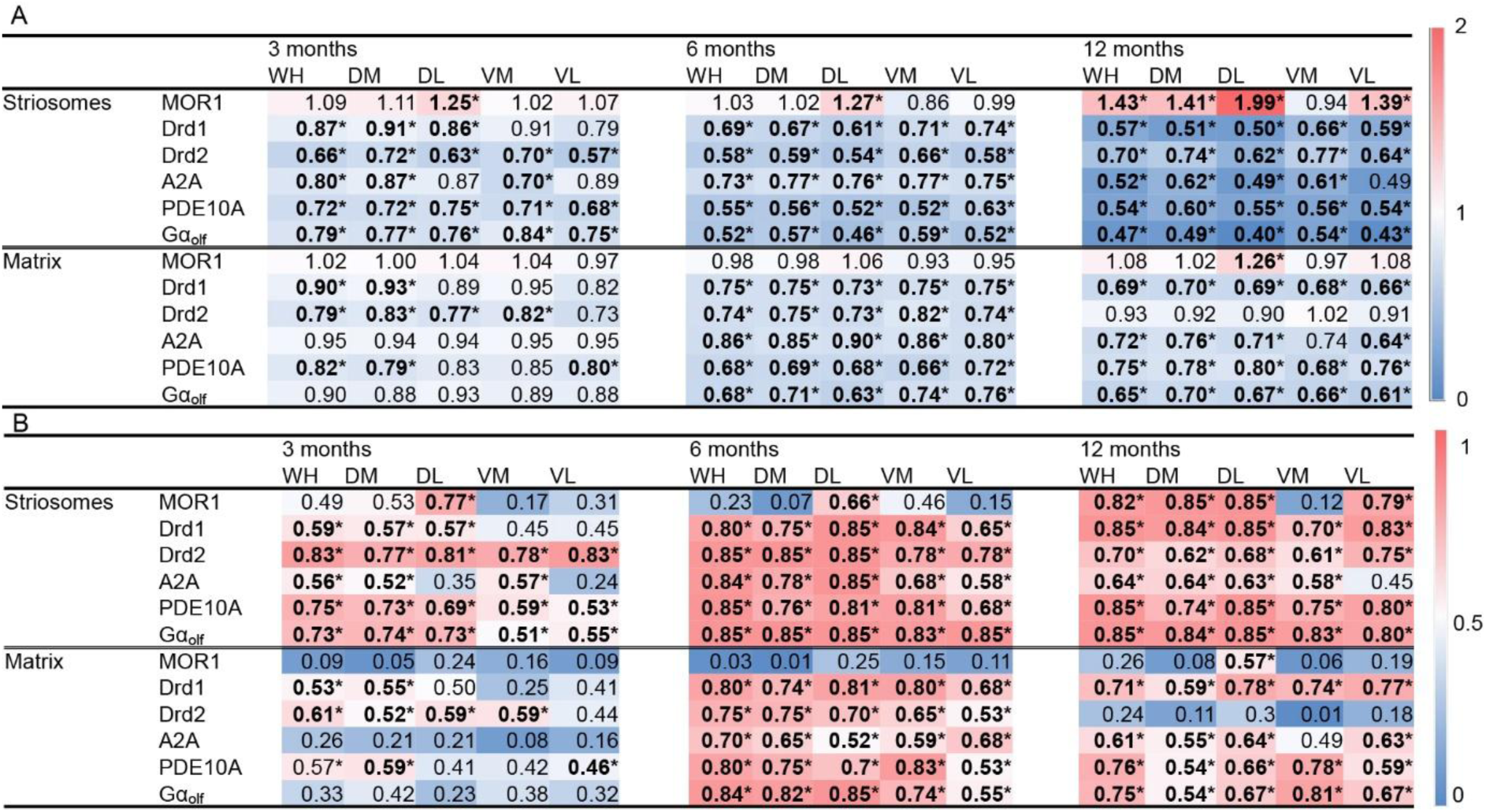
Heatmaps of changes in molecular expression in each striatal segment. The top heatmap (A) shows relative intensity, and the bottom heatmap (B) demonstrates effect size for each corresponding segment. Asterisks with values in boldface indicate statistical significance with Kruskal-Wallis tests followed by pairwise Mann-Whitney U-tests (*P < 0.005 after Bonferroni corrections).

**Figure 2:**
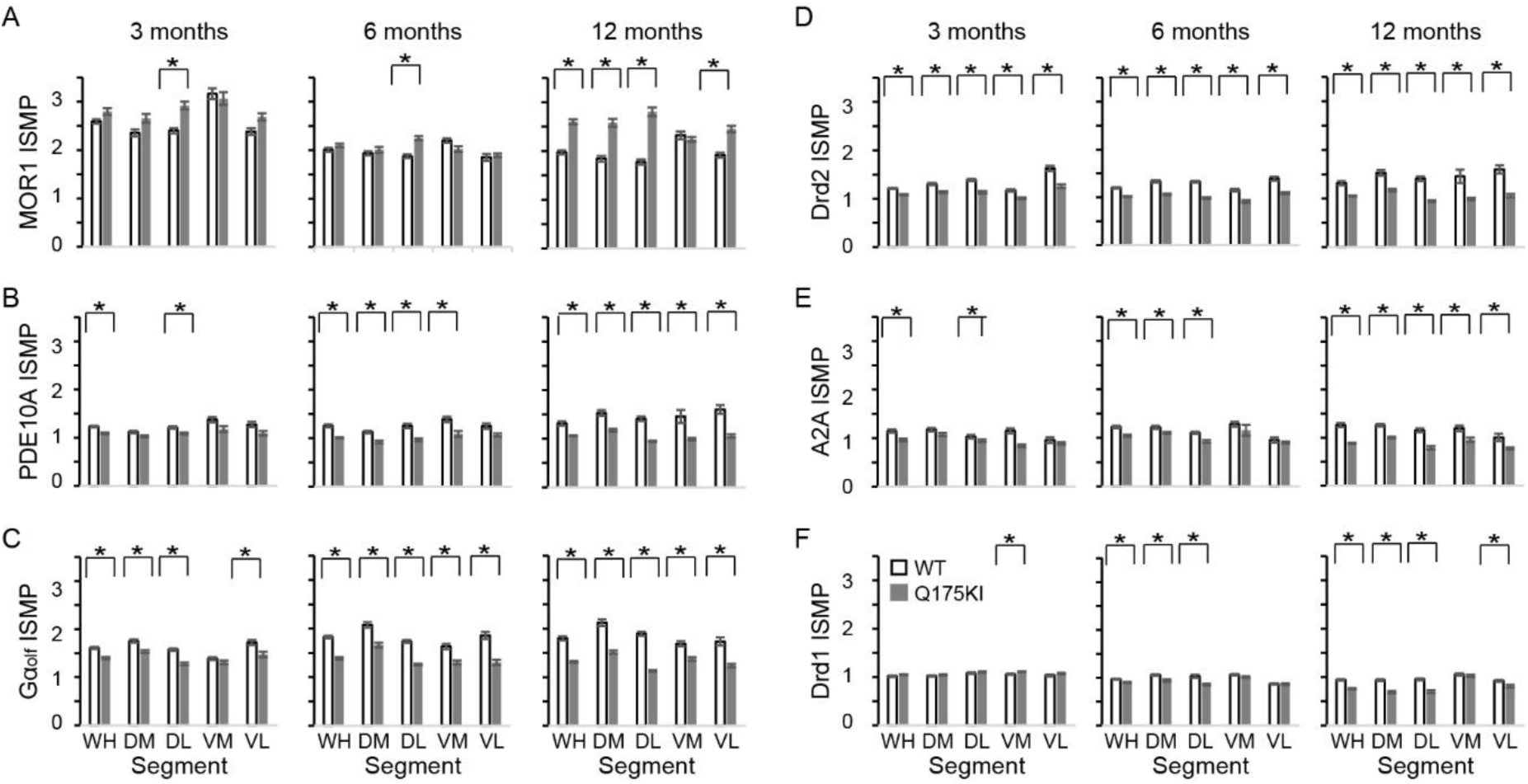
ISMP ratio of MOR1 (A), PDE10A (B), Gα_olf_ (C), Drd2 (D), A2A (E), and Drd1 (F) in WT and Q175 mice across age (3, 6, 12 months). Asterisks indicate statistical significance with Kruskal-Wallis tests followed by pairwise Mann-Whitney U-tests (*P < 0.005 after Bonferroni corrections).

### 3. PDE10A immunoreactivity over age

#### 3.1 PDE10A expression is predominant in the striosome compartment

Immunizing peptide blocking assay showed that the labeling of PDE10A was blocked when the primary antibody was preabsorbed with 0.05 µg/ml and 0.5 µg/ml PDE10A peptide antigen. At the concentration of 0.5 µg/ml, the PDE10A labeling was completely blocked and rendered equivalent to the level of negative control staining (Supplementary Figure 2). PDE10A labeling of 3-month-old naïve mice showed that PDE10A was abundant in the caudoputamen of both mice and macaque monkeys (Figure 3A-F). The striosome compartment’s immunoreactivity appeared to be slightly stronger than that of the surrounding matrix. Double immunostaining of PDE10A and MOR1 also demonstrated this slight predominance of PDE10A expression in the striosome compartment (Figure 3G-I). These visual impressions were confirmed by the densitometric analyses of 3-month-old WT mice. The PDE10A labeling was slightly but significantly stronger in striosomes than in the matrix in the WH, DL, and VM segments (n = 16, Figure 3J). The PDE10A labeling in the WH segment of WT mice was stronger in striosomes compared to matrix across 3, 6, and 12 months of age (n = 16, Figure 3K). At the level of the substantia nigra, we found massive fiber staining of PDE10A in the substantia nigra pars reticulata (SNr) (Figure 3L) but almost no immunoreactivity in the substantia nigra pars compacta (SNc) (Figure 3M, N). High magnification fields showed that some PDE10A-positive fibers ran through the SNr (Figure 3O) and eventually combined with the dendron bouquets at the SNr (Figure 3P, Q) [51,52]. These findings might indicate that PDE10A immuno-positive SPNs in the caudoputamen not only send massive axon fibers to the SNr, but likely also could influence the dopaminergic neurons in the SNc via the striosome-dendron bouquets [52].

**Figure 3:**
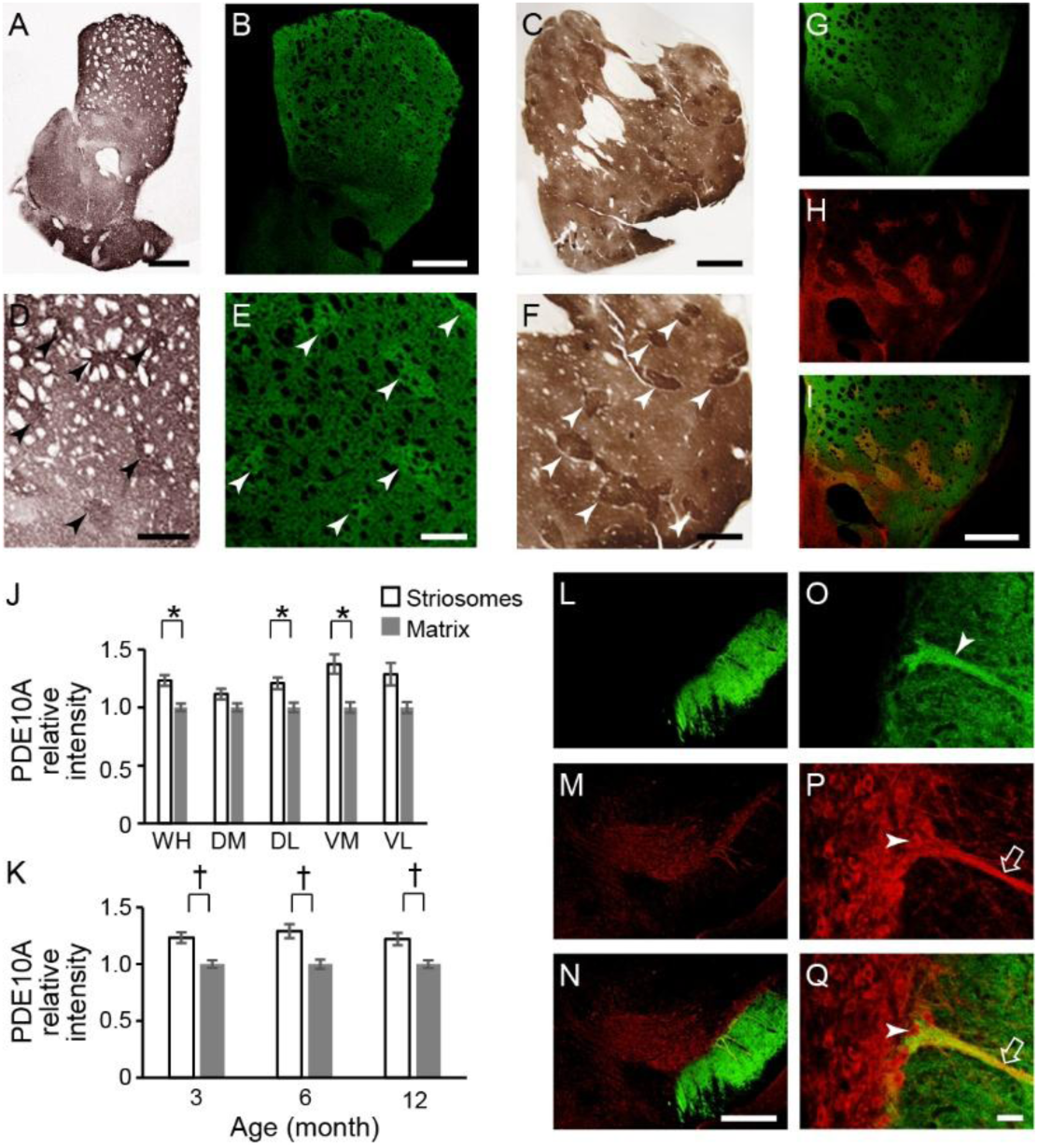
Compartmental distribution of PDE10A labeling in the caudoputamen and substantia nigra in a 3-month-old WT mouse. A-F: Microscopic images show DAB staining of mouse caudoputamen (A and D), TSA staining of mouse caudoputamen (B and E), and DAB staining of monkey caudoputamen (C and F) with low (A-C) and high (D-F) magnification. PDE10A showed stronger immunoreactivity in the dorsal caudoputamen and was slightly enriched in the striosome compartment. G-I: PDE10A labeling in the mouse caudoputamen (G), the same section double-stained with MOR1 (H), and merged image (I). J, K: Densitometric analysis showed that PDE10A immunolabeling in striosomes is significantly higher than that in matrix in the WH, DL, and VM segments of a 3-month-old WT mouse (J), and in the WH segment across all ages in WT mice (K). Kruskal-Wallis tests followed by pairwise Mann-Whitney U-tests (*P < 0.005 in J and † P < 0.0167 in K after Bonferroni corrections). L-Q: At the level of substantia nigra, PDE10A staining showed massive fiber staining in the SNr (L) with some PDE10A-positive fiber passing through substantia nigra reticulata (O; arrowhead) and directly attached to the dendron bouquet positive for tyrosine hydrogenase (P; arrow) of dopamine-containing neurons (P, Q; arrowhead). M and P show tyrosine hydrogenase staining in SNc, and N and Q show merged images. Scale bars: 200 µm (A, B, C, I, N), 100 µm (D, E, F), and 20 µm (Q).

#### 3.2 PDE10A exhibited decreased expression in Q175KI mice

We examined PDE10A expression in WT and Q175 mice across 3, 6, and 12 months of age. PDE10A immunoreactivity decreased across all ages in all segments in the striatum of the Q175KI mice except the DL and VM segments in the matrix compartment at 3 months of age (Figures 1 and 4). ISMP ratios were decreased in all but DM, VM, and VL segments at 3 months and VL segment at 6 months (Fig. 1B).

**Figure 4:**
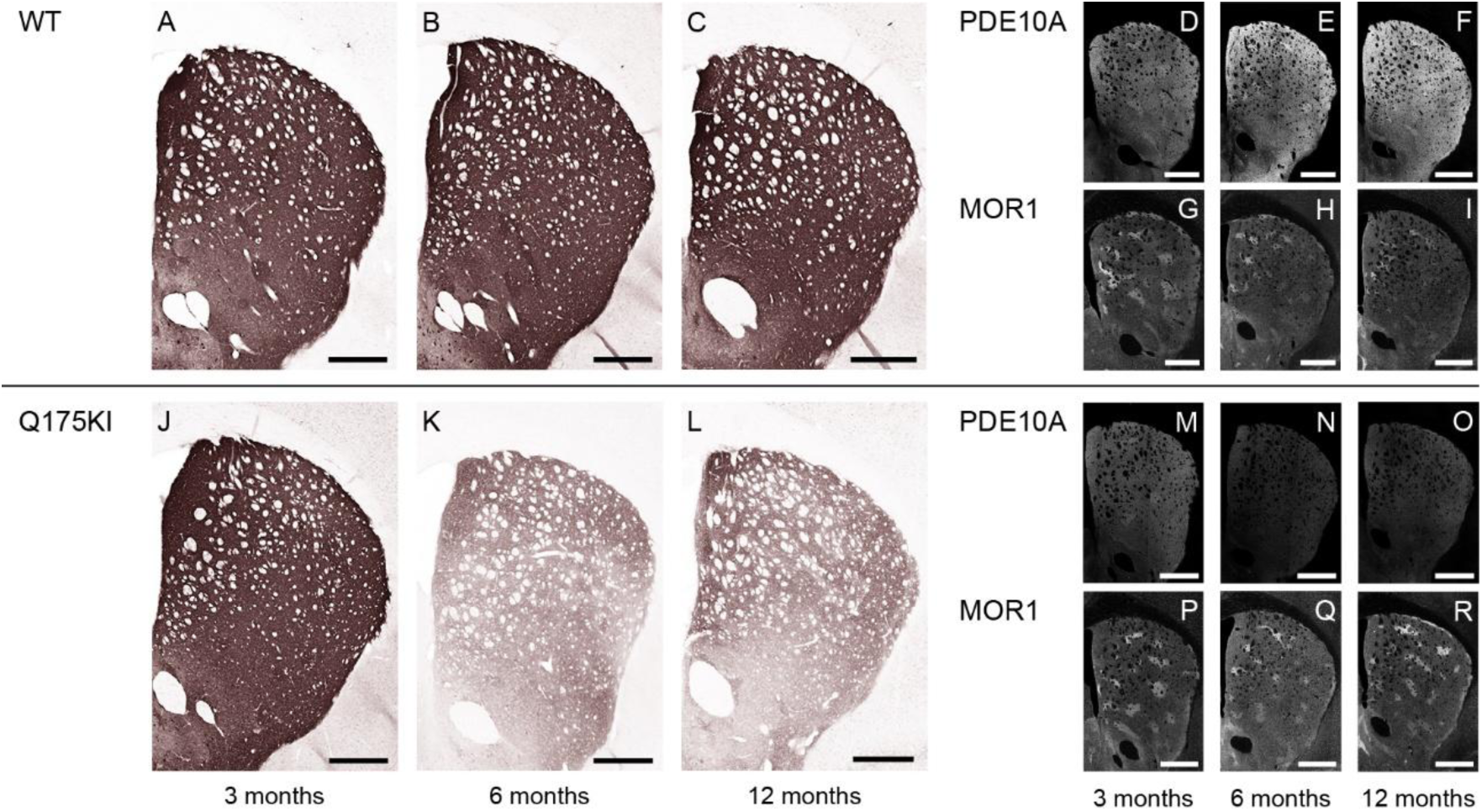
Microscopic images of PDE10A labeling at 3, 6, and 12 months. A-C, J-L: DAB-stained caudoputamen showing PDE10A in WT (A-C) and Q175KI (J-L) mice at 3, 6, and 12 months of age. D-I, M-R: TSA-stained caudoputamen showing PDE10A (D-F, M-O) double-stained with MOR1 (G-I, P-R) in WT and Q175KI mice across age groups. Scale bars: 500 µm.

### 4. Gα_olf_ and dopamine D2 receptor expression largely decreased as early as 3 months of age

Gα_olf_ expression decreased across all ages in all segments except in the matrix compartment in the 3-month Q175KI mice (Figures 1 and 5). The ISMP ratios decreased across all ages in all segments in the striatum except for the 3-month VM segment (Fig. 1C). Drd2 expression in the striosome compartment decreased in all segments of the caudoputamen across age groups. Regarding the matrix compartment, Drd2 decreased in the WH, DM, DL, and VM segments at 3 months, and in all segments at 6 months but not further at 12 months (Figures 1 and 6). ISMPs decreased at all ages in all segments (Figure 2D).

**Figure 5:**
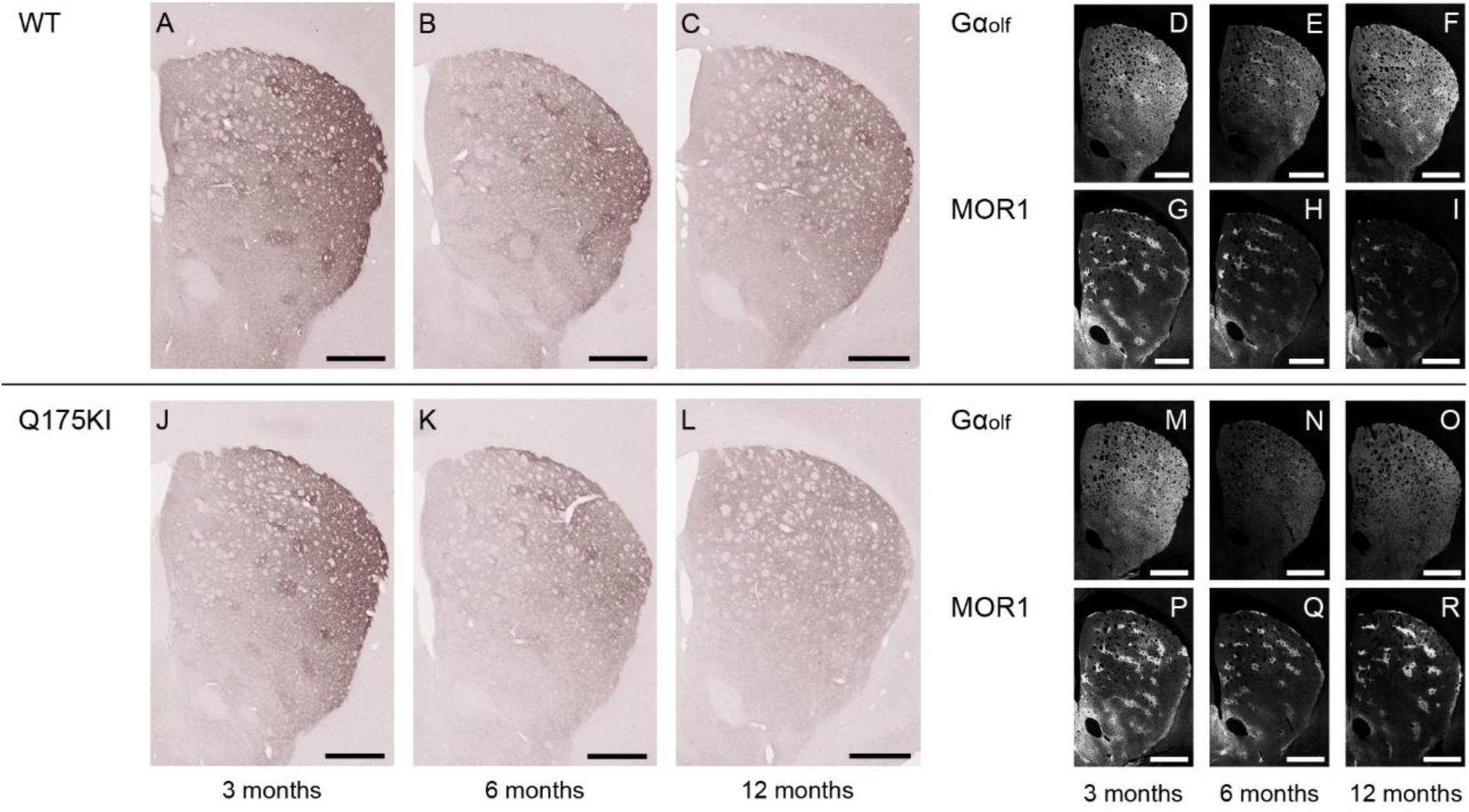
Microscopic images of Gα_olf_ labeling across 3, 6, and 12 months. A-C, J-L: DAB-stained caudoputamen showing Gα_olf_ in WT (A-C) and Q175KI (J-L) mice at 3, 6, and 12 months of age. D-I, M-R: TSA-stained caudoputamen showing Gα_olf_ (D-F, M-O) double-stained with MOR1 (G-I, P-R) in WT and Q175KI mice across age groups. Scale bars: 500 µm.

**Figure 6:**
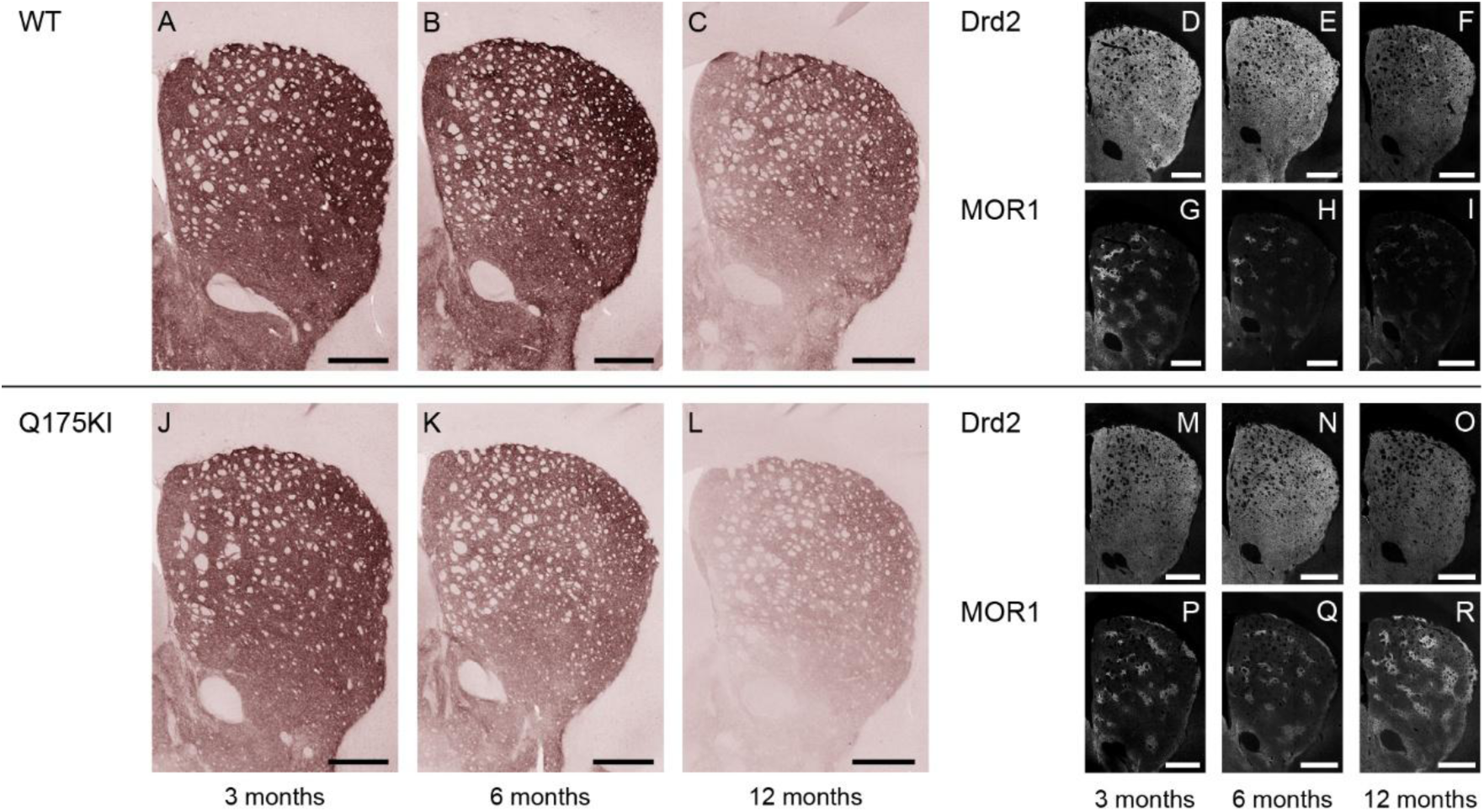
Microscopic images of Drd2 labeling across 3, 6, and 12 months. A-C, J-L: DAB-stained caudoputamen showing Drd2 in WT (A-C) and Q175KI (J-L) mice at 3, 6, and 12 months of age. D-I, M-R: TSA-stained caudoputamen showing Drd2 (D-F, M-O) double-stained with MOR1(G-I, P-R) in WT and Q175KI mice across age groups. Scale bars: 500 µm.

### 5. Adenosine 2A and dopamine D1 receptors expression exhibited relatively gradual decreases in Q175KI mice through aging

A2A expression in the striosome compartment was decreased in WH, DM, and VM segments at 3 months, and in all segments at 6 and 12 months in Q175KI mice caudoputamen (Figure 1). In the matrix compartment, A2A expression was decreased in all segments at 6 and 12 months in Q175KI mice caudoputamen. Notably, A2A decreased in striosomes in the medial (DM and VM) caudoputamen at 3 months of age and then spread to lateral (DL and VL) caudoputamen at 6 months (Figures 1 and 7). The ISMP ratio decreased in the WH and VM segments at 3 months, WH, DM and DL at 6 months, and all segments at 12 months in Q175KI mice (Figure 2E). Drd1 expression decreased across all ages in all segments except the striosome compartment of VM and VL and the matrix compartment of DL, VM and VL segments in 3-month-old Q175KI mice (Figures 1 and 8). The decreased ratio and effect size of A2A and Drd1 at 3 months of age was relatively small compared to those of PDE10A, Gα_olf_, and Drd2 (Figure 1). Drd1 ISMP increased in 3-month VM and decreased in 6-month WH, DM, DL, and in 12-month WH, DM, DL, and VL segments (Figure 2F).

**Figure 7:**
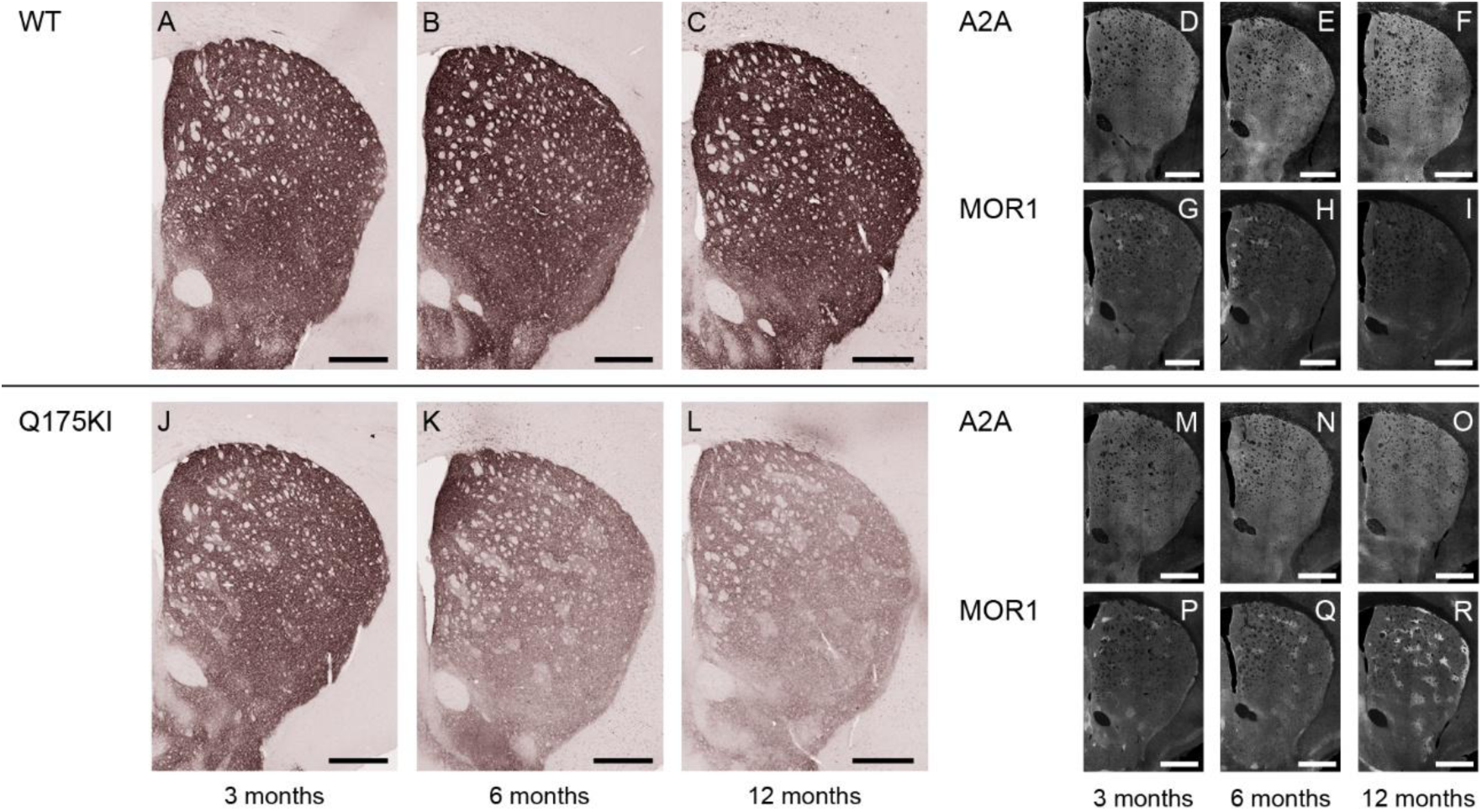
Microscopic images of A2A labeling across 3, 6, and 12 months. A-C, J-L: DAB-stained caudoputamen showing A2A in WT (A-C) and Q175KI (J-L) mice at 3, 6, and 12 months of age. D-I, M-R: TSA-stained caudoputamen showing A2A (D-F, M-O) double-stained with MOR1 (G-I, P-R) in WT and Q175KI mice across age groups. Scale bars: 500 µm.

**Figure 8:**
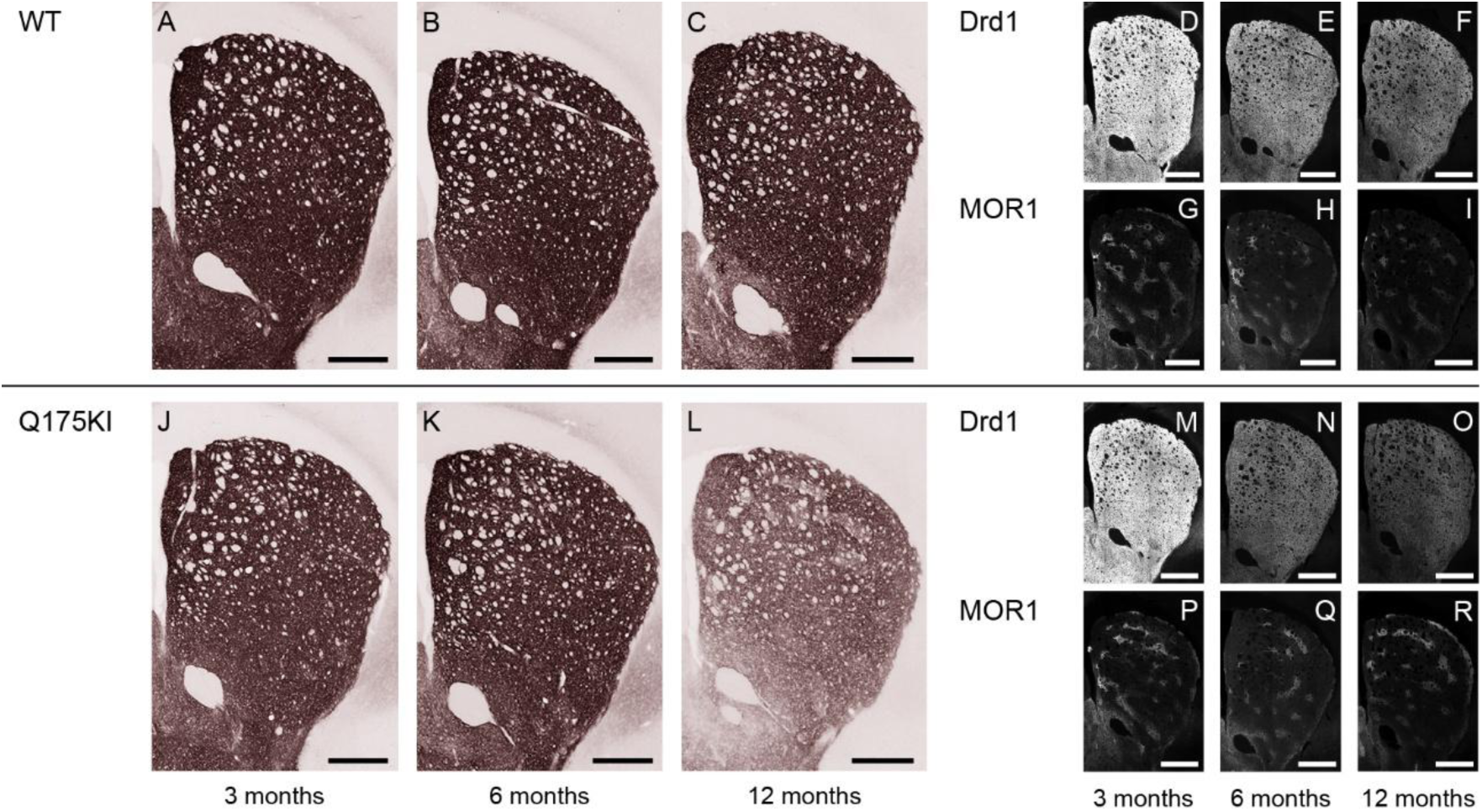
Microscopic images of Drd1 labeling across 3, 6, and 12 months. A-C, J-L: DAB-stained caudoputamen showing Drd1 in WT (A-C) and Q175KI (J-L) mice at 3, 6, and 12 months of age. D-I, M-R: TSA-stained caudoputamen showing Drd1 (D-F, M-O) double-stained with MOR1 (G-I, P-R) in WT and Q175KI mice across age groups. Scale bars: 500 µm.

## Discussion

In this study, we developed an unbiased machine learning algorithm to measure quantitatively immunohistochemically detectable signals for a panel of major receptors and second messengers across distributed sectors of the striatum of Q175KI mice and their littermate WT mice, distinguishing striosomes and matrix automatically. This improved method allowed consistent, comparable measurements to be made across different ages. We concentrated on signaling molecules affecting cAMP processing and documented here decreases of PDE10A, Gα_olf_, A2AR, Drd1, and Drd2 expressions in Q175KI mice compared to WT, detectable as early as 3 months of age (Figure 1). Their changes lasted long-term, up to 12 months of age, which indicates that downregulation of mRNA levels due to the transcriptional dysregulation evoked by mutant huntingtin protein plus the post-translation compensatory reactions induced long-lasting changes in the relative status of these cAMP-related molecules as the mice harboring the huntingtin mutation aged. In the DL segment, the so-called sensorimotor striatum, the striosome compartment was affected in the earliest stage. The imbalance between the striosome-matrix system could contribute to the appearance of movement dysfunction [50]. In addition, some of the early striosomal changes might account for mood disorders ahead of movement disorders.

### Unique status of MOR1 protein expression in HD mutants

The calculated ratios of striosome to matrix expression (ISMP values) demonstrated that the balance between striosome and matrix compartments progressively decreased in all molecules except MOR1 (Fig. 1). These decreases stood in sharp contrast to the upregulation of MOR1 expression: MOR1 ISMP ratios increased, especially at 12 months of age (Fig. 1). As seen by our results, the increase in MOR1 and the decrease in Drd1 are slow compared to those of the other molecules tested throughout the ages, indicating that alterations of these two molecules also might stem from different mechanisms than the others. The VM segment presented a different set of values, compared to the dorsal striatum, as previously noted [44]. In the VM segment, the Drd1 ISMP increased, and the Drd2 and A2A ISMP ratios decreased at 3 months of age. The Drd1 ISMP did not change at 6 and 12 months of age. MOR1 immunoreactivity and its ISMP ratios did not change throughout the 12-month age limit of the study, despite the fact that the other molecules were decreasing in this segment (Figures 1 and 2). Again, MOR1 was a standout, and furthermore, different mechanisms producing the molecular changes are likely for the dorsal and ventral striatum, a recurring theme in the neuropathology of striatum-based movement disorders.

### Clues to the etiology of observed molecular changes suggested by clinical studies

We have some clues to account for these molecular changes in the previous studies of HD. PDE10A inhibition improves HD symptoms in animal models [53–55]. Considering that PDE10A has already decreased, the positive effect of further PDE10A reduction indicates that PDE10A decrease is a compensatory reaction. As for Drd2 decrease, Drd2 antagonists are effective for humans [56–62] and HD model rats [63], indicating compensatory decreases of Drd2 in HD. In other words, a molecule that changes its expression, probably to lessen cAMP, should exist before the decrease of PDE10A and Drd2. Among the molecules examined, Gα_olf_, A2A, and Drd1 are the molecules whose downregulation could decrease the cAMP levels. A human PET study and the current study indicated that Drd1 decreases its expression relatively in the late stage (Figure 1) [14,16]. In addition, Drd1 antagonists rescue symptoms, suggesting that the Drd1 decrease might also be a compensatory reaction [64,65]. In the current study, early A2A decreases are confined to the medial caudoputamen, i.e., DM and VM segments. In addition, A2A antagonists have been reported as being neuroprotective and as alleviating symptoms in HD model mice, especially in the early stage, supporting the notion that an A2A decrease is also a compensatory reaction [66–70].

### Hypothesis: Gα_olf_ as a primary etiologic signal of abnormality in HD model mice

According to these working suggestions, Gα_olf_ might be a candidate for the primary change. Gα_olf_ couples with Drd1 and A2A and normally changes its expression depending on its usage [71]. However, in the current case, a decrease in the expression of Gα_olf_ was already observed in 3-month-old Q175KI mice, before the apparent upregulation of MOR1. As we previously hypothesized, dopamine levels parallel the upregulation in MOR1, indicating that the early Gα_olf_ decrease might not be usage-dependent due to upregulation in dopamine levels [13,44]. This timeline implies that the early Gα_olf_ decrease might be attributed to transcriptional abnormalities caused by mHtt. If translational Gα_olf_ decreases occur first, they could also induce hypersensitivity of Drd1 and A2A [71]. These hypersensitivities could promote subsequent usage-dependent consumption of Gα_olf_.

In the material analyzed here, the early striosomal changes in all segments were common in the molecules related to Drd2, as identified previously by snRNA-seq analysis [13]. Early loss of Drd2-type SPNs has also been found in HD postmortem studies [3,6,72,73]. The decrease in A2A/Gα_olf_ signaling in the striosome compartment could lead to a subsequent downregulation in Drd2/Gi signaling to compensate for the cAMP decrease. Loss of function in Drd2-type SPNs could in turn downregulate dopamine release and lead to parkinsonism [52]. To support this notion, parkinsonism is detectable early on in premanifest gene carriers up to two decades before the clinical manifestation of HD symptoms [74]. In addition, a decrease in Drd1/Gα_olf_ signal in the striosome compartment in later stages could produce upregulation of dopamine release in the striatum via striosomal pathways [52], and if so, this might induce chorea. Drd1-type SPNs in the striosome compartment are thought to have suppressive reciprocal innervation of the dopamine-containing neurons of the SNc [52]. As for the striosome-dendron bouquets, complex timing of signals occurs within the substantia nigra itself, and this is a major factor that needs study [51]. In the simplest view, downregulation in Drd1/Gα_olf_ SPNs might lead to upregulation of dopamine-containing neurons and their release of dopamine in the striatum, and thereby prompt further use-dependent Gα_olf_ decrease in Drd1-type SPNs.

### Dopamine-related changes could be accounted for by the changes in molecular signaling demonstrated here

This hypothesis could help to resolve the major unresolved question of why and how dopamine is upregulated in HD. Clinical evidence that dopamine-depleting agents such as tetrabenazine are effective in patients with HD indicates that they have excessive striatal dopamine release. Thus, dopamine upregulation might be compensated by Drd1 decrease and MOR1 increase in Drd1-type SPNs in the later stages. Normally, MOR1 in the striosome are enriched in the direct pathway Drd1-type SPNs [75–78]. This might account for the intense changes of both Drd1 and MOR1 that occur in the later stage.

### The potential role of PDE10A and changes in cAMP

In regard to clinical work, we realize that research findings on mouse models of HD models are imperfect indicators of the status of the HD striatum. They could even be misleading. There is, however, reason to consider our findings in relation not only to HD but to other movement disorders. We found PDE10A to be slightly but significantly more strongly expressed in striosomes than in matrix in neurotypical rodents and non-human primates, and found that PDE10A strikingly decreased its expression in the striosome compartment in the Q175KI mice. Childhood-onset familial chorea cases with *PDE10A* mutations and PDE10A reduction in HD are the focus of recent studies in the field of movement disorders [20–22,79,80]. As we have noted, PDE10A is a critical regulator of cAMP in both direct and indirect pathway neurons [18]. Gα_olf_ is a G-protein that couples with dopamine D1 receptors in SPNs and also with adenosine A2A receptors that co-localize with D2 receptors in SPNs [35,36]. A mutation of *GNAL,* which encodes Gα_olf_, also induces hereditary dystonia, another form of hyperkinetic movement disorder [28]. Further, Gα_olf_ is also distributed predominantly in striosomes, and its alteration in striosomes is correlated with levodopa-induced dyskinesia in Parkinson’s disease mouse models [23,36]. These coincidences suggested that dysregulation of the cAMP system in striosomal SPNs could be a key factor in the genesis of hyperkinetic movement disorders including chorea, dystonia, and levodopa-induced dyskinesia [23,81].

## Conclusions

Here we report a coordinated study demonstrating massive and chronic changes in the expression of multiple molecules in the Q175KI mouse striatum. All but MOR1 progressively decreased their expression, primarily in the striosomes, and diminished their trans-compartmental differential expressions. At the earliest stage examined, 3 months of age, which might be equivalent to the preclinical stage in human Huntington’s cases, molecules related to Drd2-type SPNs predominantly changed their expression. At the later stages (6 and 12 months of age), which might be equivalent to clinical stages of Huntington’s disease, the decrease was more evident in the molecules related Drd1-type SPNs. The imbalance of striosome-matrix cAMP-related molecules might be a key to delineating Huntington’s disease symptoms.

A significant limitation of our study is the small number of molecules analyzed due to the limited number of sections per mouse, which might not be enough to clarify the precise pathological changes. As noted in the snRNA-seq study of Q175KI mice, there is multiplexing of transcriptional changes in direct (Drd1-type) and indirect (Drd2-type) SPNs [13]. Abnormalities in the striosome compartment have further been linked to some neuropsychiatric disorders [37]. Currently, several PDE10A inhibitors are being examined as therapeutics in patients with schizophrenia [82–85]. These trials might shed light on the interaction between the cAMP system in the striosome compartment and psychiatric disease [86,87]. Our findings join these studies in suggesting that targeting altered cAMP levels in the striosome compartment might improve symptoms in patients with HD.

## Author Contributions

Conceptualization, Ryoma Morigaki and Ann Graybiel; Data curation, Ryoma Morigaki and Joji Fujikawa; Formal analysis, Ryoma Morigaki and Joji Fujikawa; Funding acquisition, Ryoma Morigaki and Ann Graybiel; Methodology, Ryoma Morigaki, Jill Crittenden and Ann Graybiel; Project administration, Ann Graybiel; Resources, Ryoma Morigaki and Tomoko Yoshida; Supervision, Jill Crittenden and Ann Graybiel; Validation, Ryoma Morigaki and Ann Graybiel; Writing – original draft, Ryoma Morigaki; Writing – review & editing, Ryoma Morigaki and Ann Graybiel. All authors have read and agreed to the published version of the manuscript.

## Funding

This work was supported by the CHDI Foundation (A-5552), the Saks-Kavanaugh Foundation, the Kristin R. Pressman and Jessica J. Pourian’13 Fund, JSPS KAKENHI Grants 17K10899 and 16KK0182, and the Beauty Life Corporation.

## Institutional Review Board Statement

Not applicable.

## Informed Consent Statement

Not applicable.

## Data Availability Statement

The original contributions presented in this study are included in the article/supplementary material. The original images and raw data can be accessed via DOI:10.5281/zenodo.15878997. Further inquiries can be directed to the corresponding author.

## Acknowledgments

We thank Christian Wüthrich, Dan Hu, Cody W. Carter, Alexander Friedman, Eric D. Nelson, Hideki Shimazu, Yasushi Takagi and Yasuo Kubota for their support and expert advice.

## Conflicts of Interest

The authors declare no conflicts of interest.

---

**Supplementary Figure 1:**
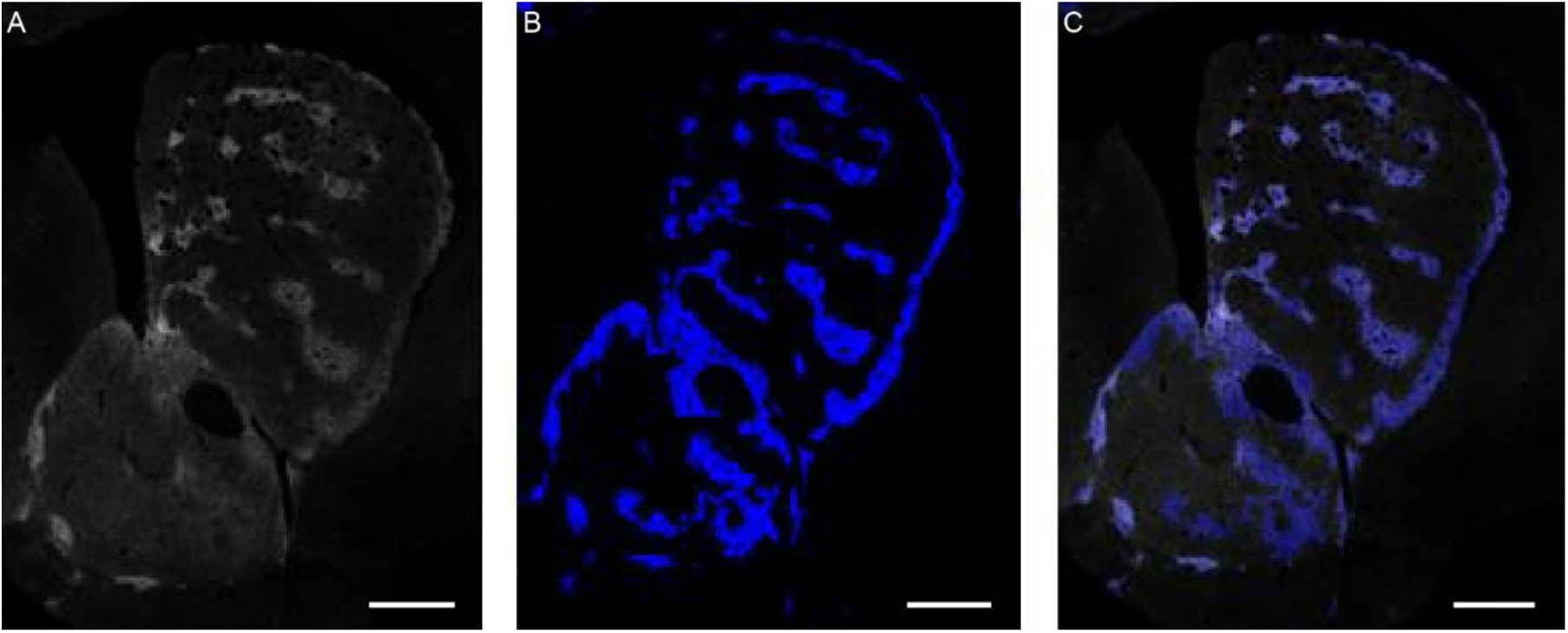
Segmentation of striosomes using U-net, demonstrated with original image (A), segmented image (B), and merged image (C). Scale bars: 500 µm.

**Supplementary Figure 2:**
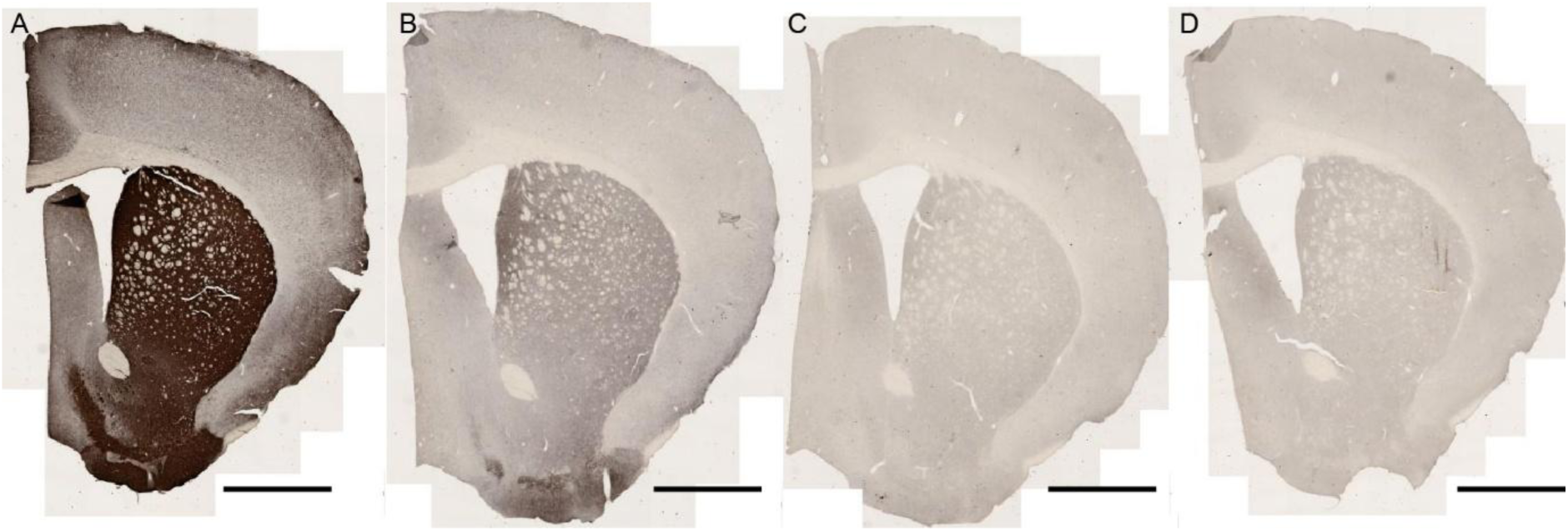
Immunizing peptide assay. In comparison to positive control staining with anti-PDE10A antibody (A), the labeling of PDE10A was blocked when the primary antibody was pre-absorbed with 0.05 µg/ml (B) and 0.5 µg/ml (C) PDE10A peptide antigen. Note that the pre-absorption with high-dose peptide antigen (C) reduced PDE10A labeling equivalent to the level of negative control staining (D). Scale bar: 1 mm.

## Notes

DOI:10.5281/zenodo.15878997

### Competing Interest Statement

The authors have declared no competing interest.

