## Supplemental Table 1 for "Molecular unbalances between striosome and matrix compartments characterize the pathogenesis of Huntington’s disease model mouse"

Supplementary Table 1: List of antibodies used in this study

| Antigen | Species raised | Company | Product Code | Dilution  DAB TSA | |
| --- | --- | --- | --- | --- | --- |
| Mu-opioid receptor | Rabbit monoclonal | Abcam, Shirley, NY | Ab134054 | 1:200 | 1:10,000 |
| K-chip | Mouse monoclonal | NeuroMab, Cambridge, MA | 75-003 | 1:2,000 | NP |
| Dopamine D1 receptor | Rat polyclonal | Sigma-Aldrich, St. Louis, MO | D2944 | 1:1,000 | 1:100,000 |
| Dopamine D2 receptor | Rabbit polyclonal | Millipore, Billerica, MA | AB5084P | 1:200 | 1:2,000 |
| Adenosine 2A receptor | Goat polyclonal | Frontier Institute, Hokkaido, Japan | A2A-Go-Af700 | 1:5,000 | 1:200,000 |
| Olfactory type G-protein α subunit | Rabbit polyclonal | Sigma-Aldrich, St. Louis, MO | SAB4501222 | 1:1,000 | 1:10,000 |
| Phosphodiesterase10A | Rabbit polyclonal | Creative Diagnostics, Shirley, NY | DPABH-03487 | 1:10,000 | 1:100,000 |
| Tyrosine hydroxylase | Mouse monoclonal | ImmunoStar, Hudson, WI | 22941 | NP | 1:20,000 |

NP: not performed.
