## Supplemental Table 2 for "Molecular unbalances between striosome and matrix compartments characterize the pathogenesis of Huntington’s disease model mouse"

Supplementary Table 2: Performance comparison of the two models of U-net

| Model | Intersection  over Union (IoU) | Precision | Recall | Dice |
| --- | --- | --- | --- | --- |
| Without augmentation | 0.689788 | 0.749693 | 0.886441 | 0.809933 |
| Augmentation with a vertical flip | 0.674448 | 0.736091 | 0.87602 | 0.797235 |
